# Network-specific synchronization of electrical slow-wave oscillations regulates sleep in *Drosophila*

**DOI:** 10.1101/542498

**Authors:** Davide Raccuglia, Sheng Huang, Anatoli Ender, M-Marcel Heim, Desiree Laber, Agustin Liotta, Stephan J Sigrist, Jörg R P Geiger, David Owald

## Abstract

Slow-wave rhythms characteristic of deep sleep oscillate in the delta band (0.5 – 4 Hz) and can be found across various brain regions in vertebrates. Across systems it is however unclear how oscillations arise and whether they are the causal functional unit steering behavior. Here, for the first time in any invertebrate, we discover sleep-relevant delta oscillations in *Drosophila.* We find that slow-wave oscillations in the sleep-regulating R2 network increase with sleep need. Optical multi-unit voltage recordings reveal that single R2 neurons get synchronized by sensory and circadian input pathways. We show that this synchronization depends on NMDA receptor (NMDARs) coincidence detector function and on an interplay of cholinergic and glutamatergic inputs setting a resonance frequency. Genetically targeting the coincidence detector function of NMDARs in R2, and thus the uncovered mechanism underlying synchronization, abolished network-specific slow-wave oscillations. It also disrupted sleep and facilitated light-induced wakening, directly establishing a causal role for slow-wave oscillations in regulating sleep and sensory gating. We therefore propose that the synchronization-based increase in oscillatory power likely represents an evolutionarily conserved, potentially ‘optimal’, strategy for constructing sleep-regulating sensory gates.

## Introduction

In vertebrates, oscillatory electrical compound patterns are associated with fundamental brain functions and specific behaviors^1-3^. Characteristic of vertebrate deep sleep are compound slow-wave oscillations in the delta band (0.5 – 4 Hz) which are thought to derive from the synchronization of neuronal activity^3^. However, how specific neural networks contribute to generating compound oscillations and whether these oscillations are the causal functional unit for sleep regulation remains unclear. This is largely due to methodological constraints, as read-outs either focus on single-cell/unit or local compound potential recordings [electroencephalograms, local field potentials] that, confined by poor spatial resolution, do not permit for dissection of multi-unit interactions ^4^.

Like vertebrates, invertebrates sleep^5-7^ and behavior selection is sensitive to an animal’s sleep need. However, in invertebrates it is unknown whether neural oscillations can gate specific behaviors, and whether an electrophysiological sleep correlate, such as slow-wave oscillations, exists or is involved in sleep regulation. Local field potential (LFP) measurements in the *Drosophila* brain indicate that the frequency of large scale compound neuronal activity is reduced during sleep^8-10^ opening up the possibility that, comparable to vertebrates, slow oscillatory activity could be involved in mediating sleep. Yet, such oscillations so far have not been identified and it remains unknown, which and how neural networks would generate slow-wave oscillations that could be crucial for sleep regulation.

We here make use of recent technological advancements in *Drosophila melanogaster* and combine targeted expression of a genetically-encoded voltage indicator (GEVI)^11-13^ with that of optogenetic actuators. This all-optical electrophysiological approach bypasses common methodological constraints allowing us to monitor multi-unit electrical patterns within a specific network and gain mechanistic insight into how sleep-relevant neural activity might be generated.

Here, we discover sleep regulating network-specific delta oscillations within the R2 network of the *Drosophila* ellipsoid body, which is situated at a crossroad involved in sleep regulation^14-16^ and multi-sensory relay for higher-order neural processing^17-19^. We demonstrate that these delta oscillations depend on multi-unit synchronization mediated through NMDA receptor (NMDAR) coincidence detection. Disrupting this synchronization and thus the emergence of compound delta oscillations disrupts sleep and alters sensory gating during sleep. We thus identify slow-wave oscillations as an electrophysiological correlate for sleep regulation in invertebrates and place these oscillatory patterns at the basis of behavior. The sleep-regulating oscillations are comparable to sleep-regulating thalamic oscillations^20-22^ as well as network-specific oscillations observed during sleep deprivation in vertebrates (local sleep)^2,23,24^. Our work demonstrates that slow-wave oscillations and sleep are fundamentally interconnected across systems, potentially representing an evolutionarily conserved strategy for network mechanisms regulating internal states and sleep.

## Results

### Sleep deprivation increases network-driven delta oscillations in the sleep-regulating R2 network

Examples of rhythmic activity patterns have previously been reported in insects^9,10^. However, their source, function and interdependence with internal states (such as sleep drive), remain largely unclear. We targeted expression of the GEVI ArcLight specifically to R2 neurons in the *Drosophila* brain. This defined network of 10 cells per hemisphere (**Fig. 1a**) resides within the ellipsoid body and is involved in sleep regulation^14-16^ and multi-sensory relay^17-19^.

**Figure 1:**
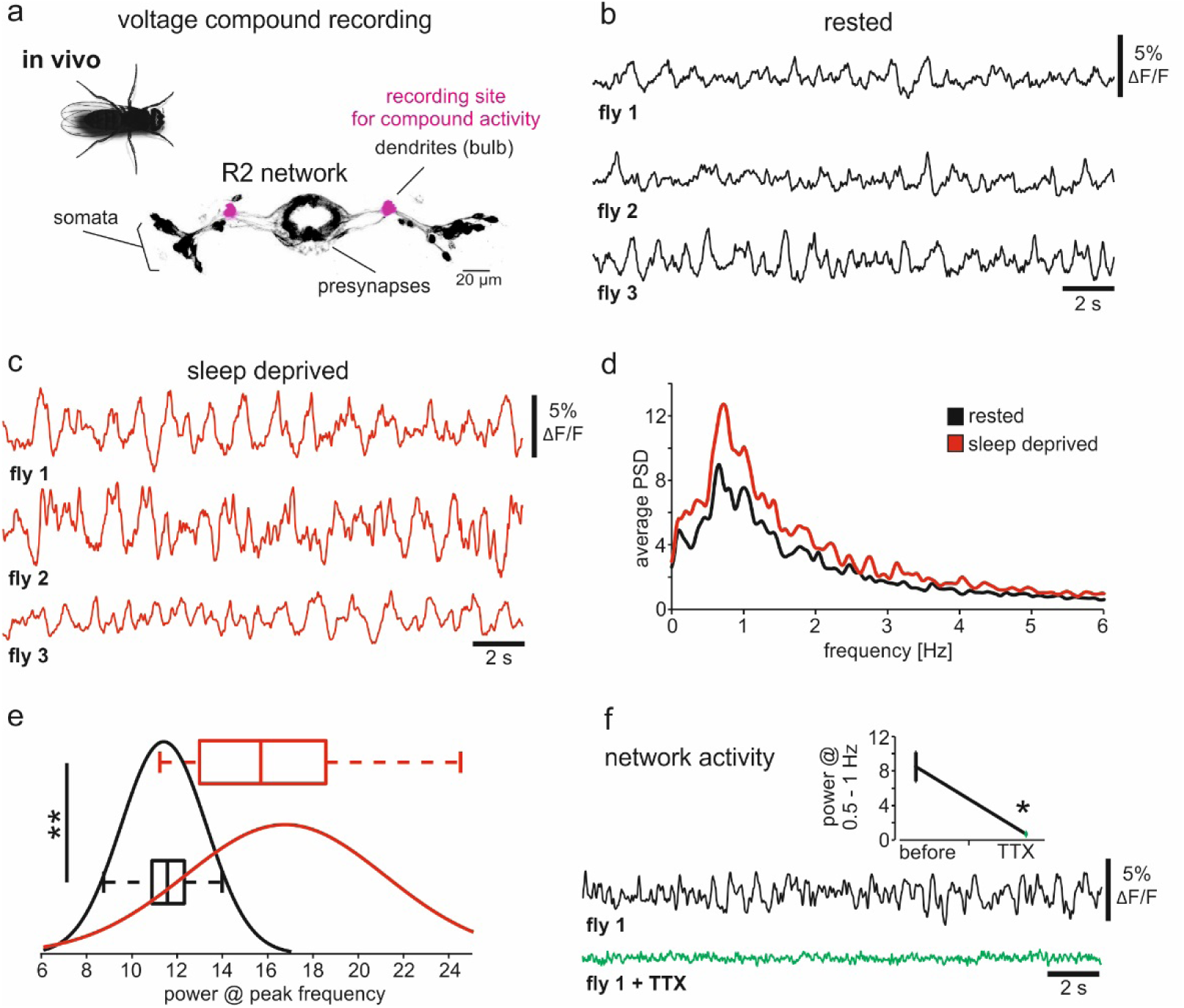
Network-driven delta oscillations in R2 neurons are sensitive to sleep need. **a,** Dendritic recording sites of R2 neurons *in vivo* (scheme). **b, c,** *In vivo* electrical compound oscillations in R2 neurons (R88F06-Gal4>UAS-ArcLight) of rested (black) and sleep-deprived flies (red). **d**) Average power spectra of voltage recordings (N=10). **e,** Distribution of oscillatory power at peak frequencies. N=10; Mann-Whitney Test, p**=0.003. **f,** TTX abolishes oscillatory patterns. N=3; paired T-Test, p*<0.05, mean± sem.

*In vivo* recordings of the dendritic processes (bulb) of R2 neurons (**Fig. 1a**) identified electrical compound activity (**Fig. 1b**) that oscillated at delta-band frequencies between 0.5-1.5 Hz (**Fig. 1b, d**) in rested flies. We sleep-deprived flies (see methods) to investigate whether these network-specific oscillations were sensitive to their internal state (**Fig. 1c**). While frequencies were unchanged (**Fig. 1d**), the power of up- and down-states was significantly increased and distributed discretely (**Fig. 1e**). Bath-applying the voltage-gated Na^+^-channel blocker TTX fully abolished electrical activity (**Fig. 1f**), indicating that delta-band oscillations require network activity. Thus, we here discovered network-specific delta-band oscillations sensitive to sleep need in invertebrates. These are comparable to brain region-specific slow-wave oscillations characteristic of ‘local sleep’ phenomena in vertebrates^2,24^. Slow-wave oscillations may therefore represent an evolutionarily conserved ‘optimal’ strategy, to convey the network state of sleepiness.

### Multicellular optical electrophysiology reveals how NMDAR-dependent single-unit activity shapes compound oscillations

To investigate the mechanisms involved in generating delta oscillations in R2 neurons, we turned to a whole-mount explant brain preparation. Consistent with the absence of external cues, compound oscillatory activity was largely reduced *ex vivo* (**Supplementary Fig. 1a**), permitting us to dissect the individual components shaping electrical network activity.

Importantly, reminiscent of mammalian preparations^25^, delta-band oscillations were restored (**Fig. 2a, b**) by increasing network activity through lowering extracellular Mg^2+^. Indeed, frequency peaks measured at low [Mg^2+^]_e_ (5 mM) *ex vivo* fully overlapped with those measured *in vivo* (**Fig. 2c**) at high [Mg^2+^]_e_ (20 mM, physiological range^26,27^ in *Drosophila*, see **Supplementary Fig. 1b**). The *ex vivo* recording configuration allowed us to measure individual somata simultaneously providing readout from multiple single-units. Individual R2 neurons showed oscillatory activity with peak frequencies similar to the compound signal, suggesting that overlap of electrical patterns generates compound oscillations (**Fig. 2d, e**).

**Figure 2:**
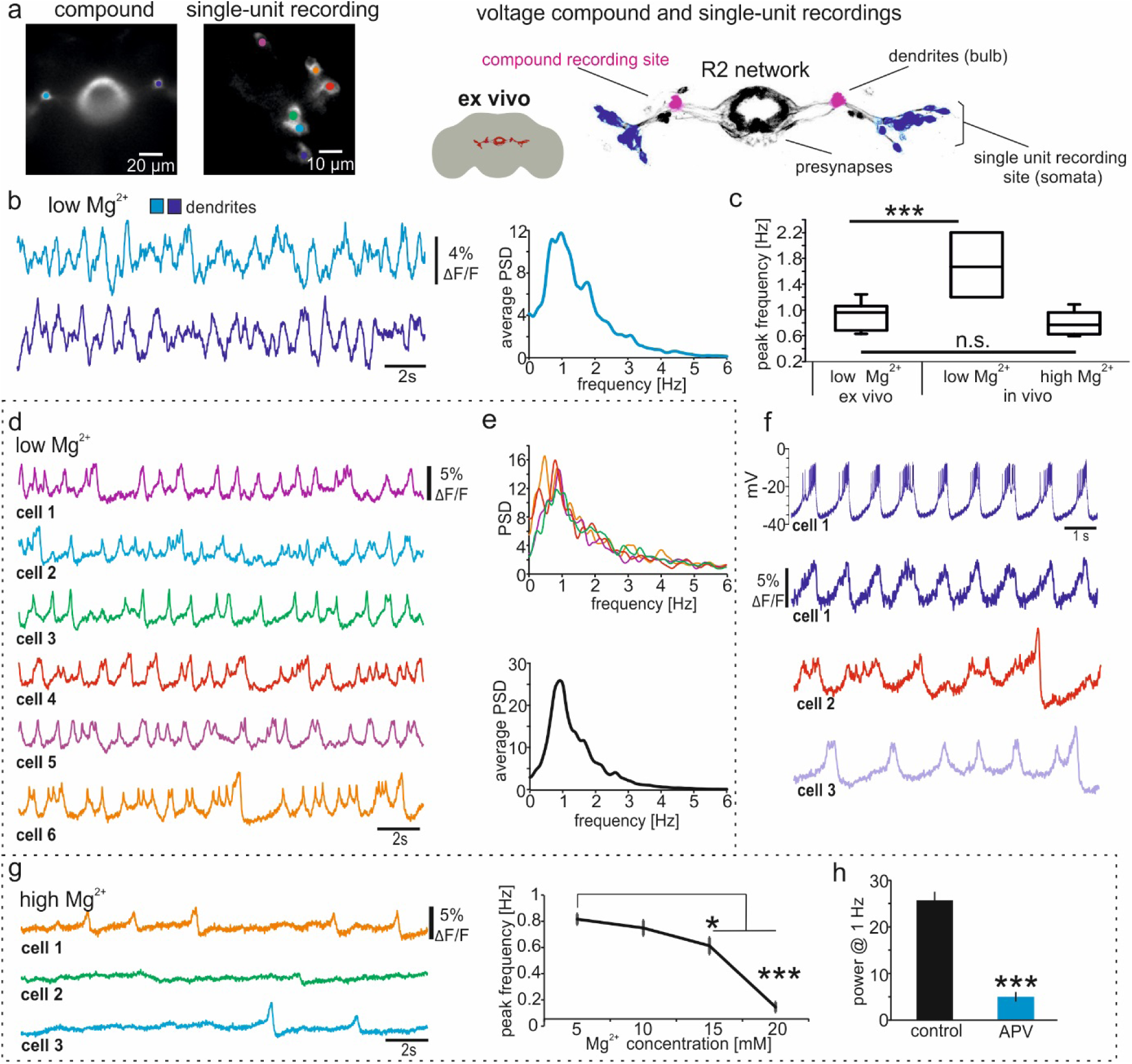
NMDAR-dependent single-unit oscillations shape compound activity. **a,** Wide-field images of R88F06-Gal4>UAS-ArcLight. Areas of interest are indicated for compound and single-unit recordings (left). Recording sites are indicated schematically (right). **b,** *Ex vivo* compound delta oscillations recorded at low (5mM) [Mg^2+^]_e_, power spectrum (right). **c,** Peak frequency distribution *ex vivo* at low [Mg^2+^]_e_ strongly resembles *in vivo* compound activity at high (20mM) [Mg^2+^]_e_. At low [Mg^2+^]_e_ *in vivo* compound activity is significantly higher. N=8-18; one-way ANOVA followed by Bonferroni, p***<0.001. **d,** Multi-cellular voltage imaging of individual R2 neurons. **e,** Power spectrum of several R2 neurons (shown in **d**). Average spectrum in lower panel (N=90, >8 flies). **f,** Multi-cellular optical and single-cell patch-clamp recording. Cell 1 is simultaneously recorded optically and via the patch pipette (blue), while two other R2 neurons are recorded optically. **g,** Electrical activity of single R2 neurons at 20mM [Mg^2+^]_e_. Frequency depends on [Mg^2+^]_e_ (right panel). N=41-48 neurons; one-way ANOVA followed by Bonferroni, 5mM [Mg^2+^]_e_ was used as control, p*=0.028, p***<0.001, mean± sem. **h,** APV (100μM) abolishes oscillations at 1Hz. N=34-45; Mann-Whitney Test, p***<0.001, mean± sem.

Combined whole-cell patch-clamp and voltage imaging experiments calibrated the observed single-unit membrane potential fluctuations to around 26 +/-7 mV. Importantly, depolarization phases were decorated with spikes, confirming that electrical patterns represented up-states, followed by down-states marked by neuronal silence (**Fig. 2f**). Single-unit activity was also sensitive to [Mg^2+^]_e_ and peak frequencies dropped close to the reported concentrations needed for a full Mg^2+^-block of *Drosophila* NMDA receptors^28^ (**Fig. 2g**). NMDARs are coincidence detectors and activation requires simultaneous ligand-binding and membrane depolarization to remove Mg^2+^-ions blocking the channel pore. To test whether NMDARs modulated oscillatory activity, we applied the selective NMDAR antagonists APV or MK-801 and found single-unit oscillations to be abolished in both cases (**Fig. 2h and Supplementary Fig. 1c**). Thus, single-unit delta-band oscillations require network activity potentially generated by NMDAR-mediated signaling. Interestingly, sleep deprivation leads to an up-regulation of NMDAR transcripts in R2 neurons^16^, opening the possibility that increased sensitivity to input pathways could be involved in shaping delta-band oscillations.

### R2 network-specific multi-unit synchronization generates delta-band compound oscillations

To investigate, whether input pathways generated or shaped delta-oscillations, we expressed the red light activatable channelrhodopsin CsChrimson in AOTU (anterior optical tubercle) neurons that provide direct sensory and circadian input to R2^14,17,19^. Compared to previous experiments (**Fig. 2**) baseline frequencies were slightly increased, probably due to low baseline activation of CsChrimson during imaging as indicated by control experiments using flies which were not fed with retinal (**Supplementary Fig. 3d**). Activation of the AOTU reinstated delta oscillations at high [Mg^2+^]_e_ and single units showed a strong reversible increase in power (**Fig. 3a, b**) reflecting increased voltage deflections of up- and down-states. Strikingly, we observed a reduction of the phase-lag and increased correlation of up-states between individual units (**Fig. 3c**). This demonstrates that activity transmitted via the AOTU can synchronize R2 neurons. Moreover, both the increase in single-unit power and multi-unit synchronization are means for increasing network compound power, as observed after sleep-deprivation (**Fig. 1**).

**Figure 3:**
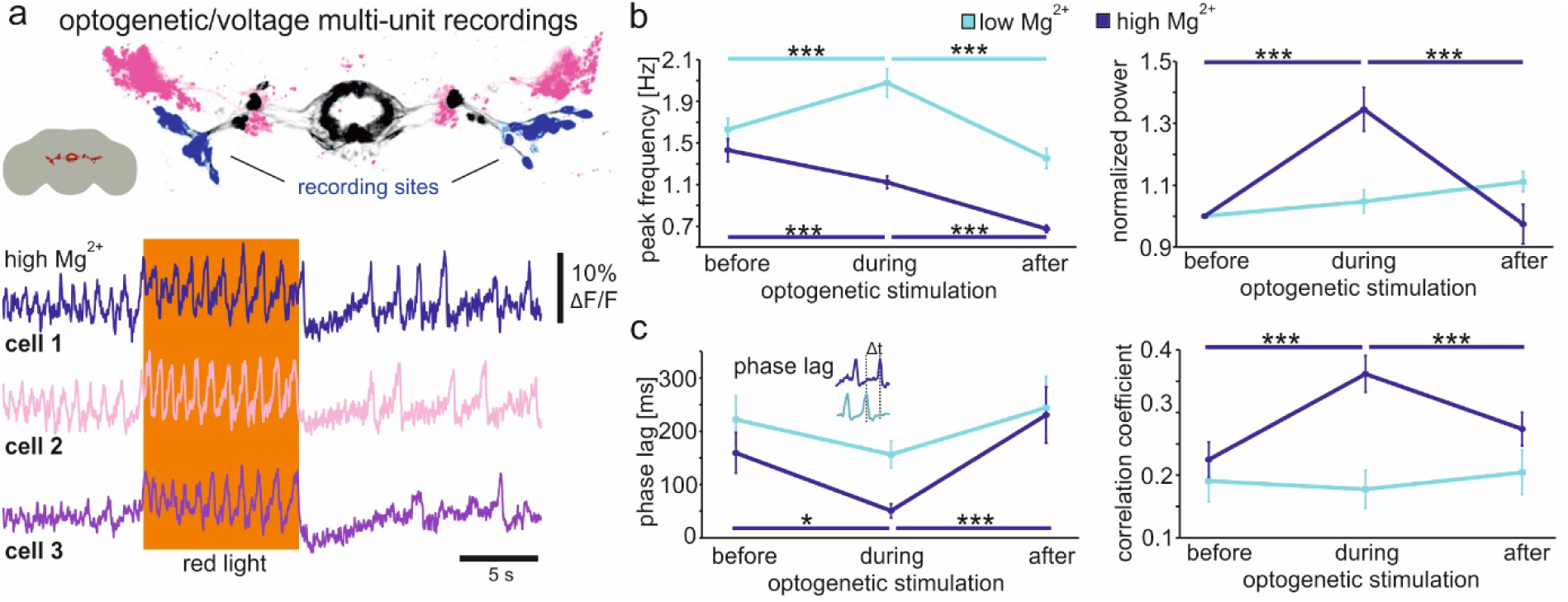
Coherent AOTU activation can synchronize R2 electrical patterns. **a,** Schematic overview of recording sites (somata blue, R88F06-Gal4). AOTU (R76B06-LexA) shown in pink. R2 neurons show changes in electrical activity during optogenetic activation of AOTU. **b,** Peak frequency and power, **c,** phase lag between up states and correlation coefficient between single units at low (5mM) [Mg^2+^]_e_ and high (20mM) [Mg^2+^]_e_ during optogenetic activation. N=51 for low, N=53 for high, at least 8 flies; One-way ANOVA on repeated measures, followed by Bonferroni, p*<0.05, p***<0.001, mean± sem.

No oscillatory activity was detected within the AOTU (**Supplementary Fig. 2**), confirming that oscillations are generated at the level of R2 neurons in response to network input. Importantly, at low [Mg^2+^]_e_, optogenetic activation of the AOTU increased oscillatory frequency, but did not change single-unit power or multi-unit synchrony (**Fig. 3b, c and Supplementary 3a, b**). Moreover, removing NMDARs ‘from the equation’ by applying APV completely prevented AOTU-induced activity in R2 units (**Supplementary Fig. 3c**). Therefore, NMDARs are required for oscillatory activity *per se*, while [Mg^2+^]_e_ levels required for the NMDAR Mg^2+^-block were decisive as to whether the network stimulation would increase oscillatory frequency or lead to multi-unit synchrony.

### Bidirectional frequency modulation via excitatory cholinergic and inhibitory glutamatergic input

NMDARs typically work in concert with other ligand-gated ion channels for coincidence detection ^29,30^. Pharmacological block of nicotinic acetylcholine receptors (nAChR) strongly reduced the frequency of baseline oscillations (**Fig. 4a, b**), indicating that cholinergic input could provide the ‘second’ signal. Cholinergic input was required at the level of R2 and not elsewhere in the circuitry, as targeting RNA interference against the evolutionarily conserved α7 subunit of nAChR in R2 also reduced oscillatory frequency (**Fig. 4a, b**). Moreover, when focally applying acetylcholine, we noticed a fast transient activation of the R2 network (*not shown*) and several consecutive puffs induced oscillatory activity. Importantly, activity induction was sensitive to the competitive NMDAR blocker MK-801 (**Fig. 4c**). Thus, acetylcholine can provide the coincident signal to ‘unblock’ NMDARs required for synchronization of single units at the basis of delta band oscillations.

**Figure 4:**
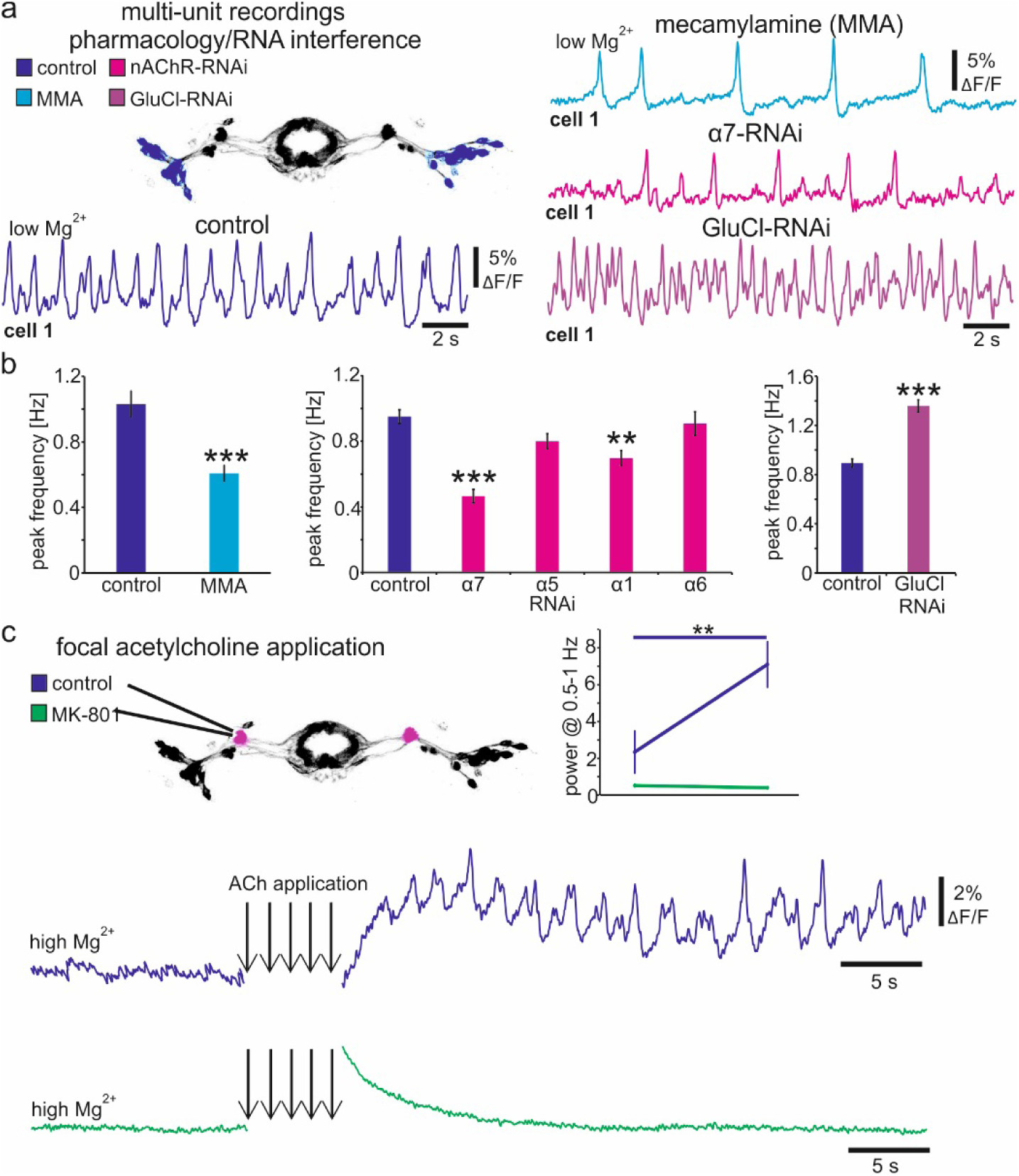
Frequency and power of R2 delta oscillations are defined by cholinergic and glutamatergic synaptic inputs. **a, b,** Multi-unit recordings after applying mecamylamine (MMA) or when cell-specifically (R88F06-Gal4) knocking-down nicotinic α subunits (N=49-66, at least 5 flies; one-way ANOVA followed by Bonferroni, p**=0.002, p***<0.001) or GluCl receptor (N=34-67 from at least 4 flies; one-way ANOVA followed by Bonferroni, p***<0.001). **c,** Induction of oscillations by consecutive focal application of acetylcholine (arrows) is abolished when applying MK-801. N=5; paired T-Test, p**<0.01. Bar graphs: mean± sem.

Interestingly, down-regulation of glutamate-gated chloride (GluCl) conductance increased the frequency of oscillations (**Fig. 4a, b**). We therefore propose that the interplay between glutamatergic and cholinergic signals sets a resonance frequency that permits temporal synchronization of single units. Indeed, our experiments indicate that, only if the frequency is kept constant around 1 Hz (or even reduced, **Fig. 3a, b**), multi-unit synchronization is induced.

### NMDAR coincidence detection specific to R2 neurons is crucial for generating sleep regulating delta oscillations

Our results indicate that NMDAR and more specifically the NMDAR Mg^2+^-block is crucially involved in generating R2-specific compound oscillations (**Fig. 2h, 3, 4c**). To test whether *in vivo* oscillatory activity in R2 neurons was directly governed by the NMDAR Mg^2+^-block we used cell-specific expression of a Mg^2+^-insensitive NMDAR subunit 1 mutant (N631Q / NMDAR^Mg-/-^)^28^. This subunit forms heteromers with the endogenous NMDAR subunit 2, generating NMDARs that are no longer blocked by Mg^2+^, and therefore lose their coincidence detector function^28^. *In vivo* whole-cell patch-clamp recordings of R2 neurons expressing NMDAR^Mg-/-^ showed tonic firing of individual neurons to be increased (**Fig. 5a-c**), resulting in aberrant compound oscillations (**Fig. 5d, e**). Interestingly, we also observed abolished compound oscillations when down-regulating NMDARs with RNA interference specifically at the level of R2 neurons (**Supplementary Fig. 4**), confirming that it indeed is the cooperativity of single units that allows for multi-unit synchronization. Importantly, expression of NMDAR^Mg-/-^ in R2 now provided us with the tools to directly test for causality of multi-unit synchronization in regulating sleep behavior.

**Figure 5:**
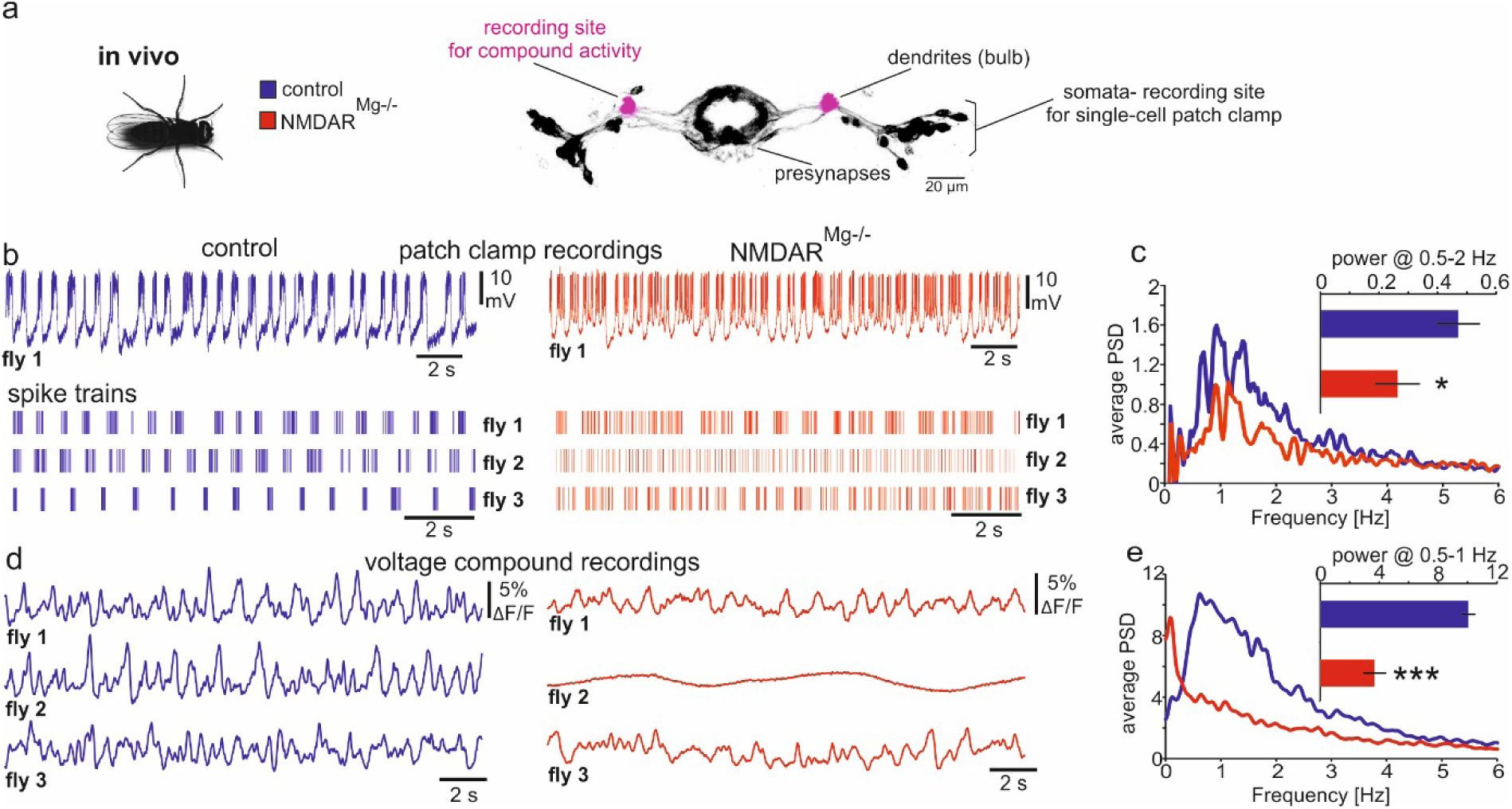
NMDAR coincidence detection in R2 neurons is crucial for generating *in vivo* compound delta oscillations. **a,** Schematic indicating recording sites. **b,** Single-cell *in vivo* patch-clamp recordings of individual R2 neurons (R88F06-Gal4) expressing UAS-dNR1 wt as controls (blue) or UAS-dNR1 N631Q (NMDAR^Mg-/-^, red). Controls show regular bursting patterns, NMDAR^Mg-/-^ leads to tonic firing. **c,** Power of delta band oscillations is reduced for NMDAR^Mg-/-^. N=4 from 4 flies; unpaired T-Test, p*=0.048. **d,** *In vivo* compound activity (optical recordings) of R2 neurons expressing NMDAR^Mg-/-^. **e,** Power spectrum and power of compound oscillations between 0.5 and 1 Hz. N=9-11; Mann-Whitney Test, p***<0.001, mean± sem.

We monitored sleep patterns of flies that expressed NMDAR^Mg-/-^ at the level of R2. Flies slept significantly less in total compared to controls (**Fig. 6a, b**) and the number of sleep episodes was increased (**Fig. 6d**), while sleep episode duration was decreased (**Fig. 6e**) leading to a largely fragmented sleep pattern (**Fig. 6c**). Moreover, sleep latency was increased (**Fig. 6f**) and activity counts during wake phases were reduced rather than increased (**Fig. 6g**), indicating that the underlying change in sleep pattern was specifically due to altered sleep regulation. Thus, flies no longer capable of multi-unit synchronization in R2 neurons woke up more frequently and it took them longer to fall asleep. This directly links R2 delta oscillations to the generation of adequate sleep drive for both the induction and the maintenance of sleep.

**Figure 6:**
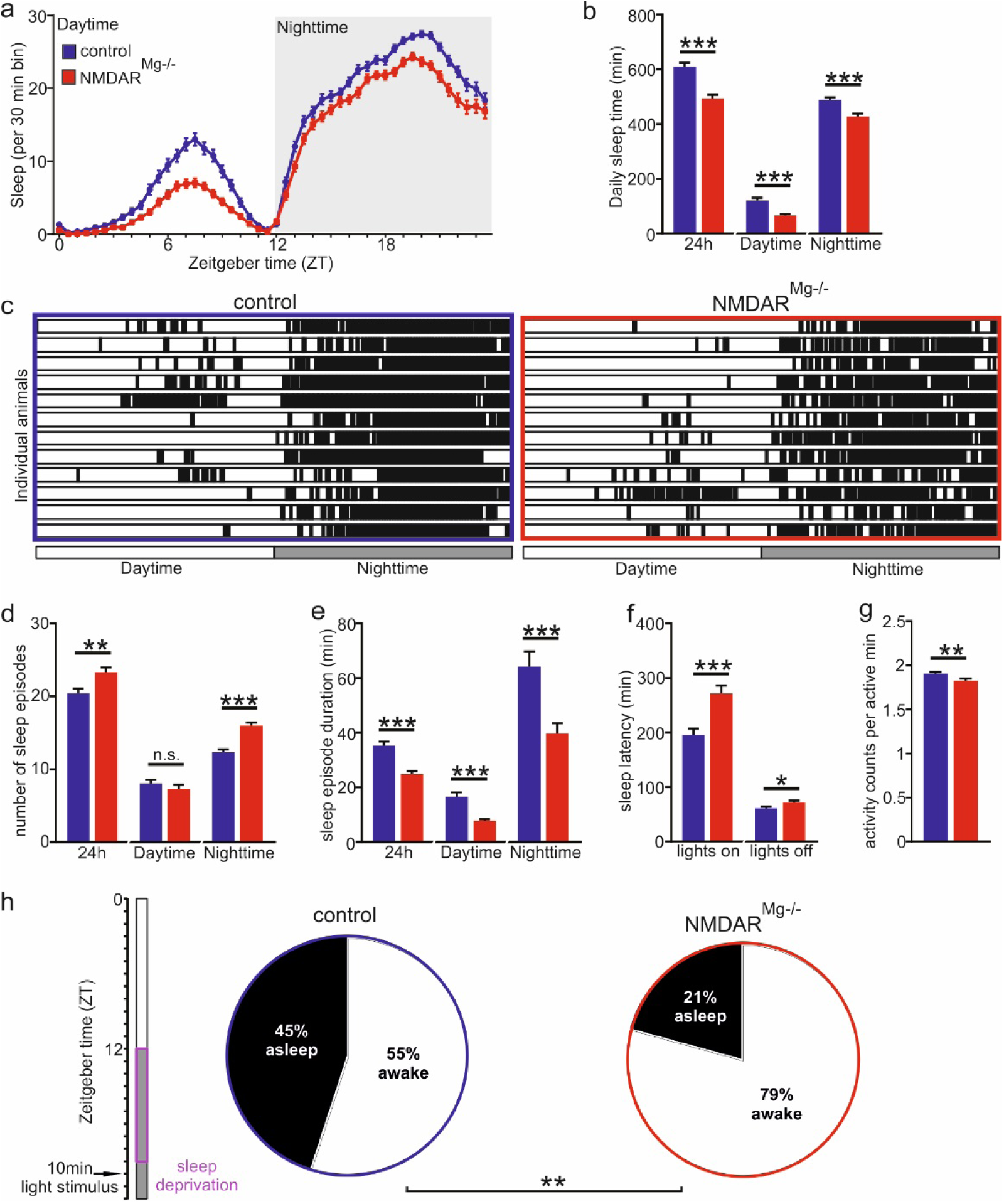
Loss of delta oscillations in R2 disrupts sleep. **a,** Sleep counts for flies expressing NMDAR^Mg-/-^ (red, UAS-dNR1 N631Q) in R2 neurons (R58H05-Gal4) and controls (blue, UAS-dNR1 wt). **b,** Expressing NMDAR^Mg-/-^ reduces average sleep amount. N=127; unpaired T-Test, p***<0.001. **c,** Sleep pattern of individual flies showing periods of wakefulness (white bars) and sleep (black bars). Expressing NMDAR^Mg-/-^ **d,** increases number of sleep episodes and **e,** reduces sleep episode duration leading to heavily fragmented sleep (N=127; unpaired T-Test, p**=0.002, p***<0.001) as well as **f,** an increase in sleep latency **g,** and a reduction in general locomotor activity (N=127; unpaired T-Test, p*=0.02, p**=0.008, p***<0.001). **h,** Expression of NMDAR^Mg-/-^ in R2 neurons (R58H05-Gal4) increases the probability of waking up when exposed to light following 9 hours of sleep deprivation. N=53-60; unpaired T-Test, p**=0.006, see methods. Bar graphs: mean± sem.

If multi-unit synchronization at the basis of sleep regulation represented an evolutionarily conserved mechanism, we reasoned that sensory-filtering, sleep maintenance and wakening should be interconnected, potentially analogous to the function of thalamic nuclei in vertebrates^31^. We sleep-deprived flies that expressed NMDAR^Mg-/-^ in R2 neurons, let the flies go to sleep and tested whether they could be woken up by visual stimuli (**Fig. 6h**). Indeed, the wake-up threshold was lower in the mutant and a significantly larger fraction of flies was wakened (**Fig. 6h**) compared to controls. Thus, delta oscillations in R2 neurons not only regulate sleep drive, but also the gating of stimulus-triggered wakening, following an evolutionarily conserved strategy.

## Discussion

In vertebrates, sleep and sleepiness are thought to be tightly interlinked with the synchronization of neuronal activity, resulting in increased compound slow-wave oscillations^2,3^. However, it remains unknown, whether all animals that sleep, including invertebrates, show network-specific slow-wave oscillations involved in sleep regulation. Conserved over evolution, synchronization could represent an ‘optimal’ strategy for sleep regulation. We here, in an invertebrate model system, uncover a NMDAR coincidence-detector based mechanism gating network-specific and sleep-relevant neuronal synchronization of delta wave activity.

Our data suggest that compound delta oscillations specific to the sleep-regulating R2-network are generated by circadian information mediating coherent synaptic output via the AOTU. We show that optogenetic activation of the AOTU increases single-unit power and synchronizes R2 neurons (**Fig. 3**), resulting in an increase of the power of compound delta oscillations (**Fig. 1**) and thus internal sleep drive. This is consistent with thermogenetic activation of the AOTU increasing total amount of sleep in flies^17^. Generating coherent AOTU output could be mediated by sleep-modulating DN1 circadian clock neurons^32^, which form direct connections with the AOTU and are likely to modulate oscillatory activity in R2 neurons^14^.

The R2 network also receives excitatory input from Helicon cells^15^, which could be a potential source of cholinergic input. We here provide evidence that nAChRs act as prime candidates to provide concurrent depolarization required for NMDAR coincidence detection in R2 neurons (**Fig. 4**). Helicon cells also receive visual information and are part of a recurrent circuit mediating homeostatic sleep regulation^15,33^. Thus, R2 oscillatory activity is likely regulated via a complex interplay of sensory input^19^, circadian rhythms^14,17^ and homeostatic sleep regulation^15^.

Our experiments further indicate that additional inhibitory input via glutamate-gated chloride channels antagonizes excitatory drives to prevent an ‘overshoot’ of frequencies, potentially defining the time-window for recurrent input (**Fig. 4**). We therefore propose that the interplay between glutamatergic and cholinergic signals sets a resonance frequency that permits temporal synchronization of single units to define the power and frequency of oscillatory activity and thus the strength of sleep drive. Of note, following AOTU-mediated synchronization, a prolonged hyperpolarization becomes apparent (**Fig. 3**), reminiscent of a *Bereitschaftspotential,* that typically can be observed following highly synchronized states^4^.

In R2 neurons expression levels of NMDAR had previously been associated with regulating sleep drive^16^. However, the mechanistic involvement of NMDAR remained elusive. We here demonstrate that NMDAR coincidence detection gates neuronal synchronization of delta wave activity within the R2 network to increase the power of sleep-relevant compound oscillations (**Fig. 3, 5**). Mg^2+^ block-deficient NMDAR in R2 neurons led to tonic firing patterns in single R2 neurons (**Fig. 5b**), which reduced delta oscillations at the compound level (**Fig. 5d, e**). This demonstrates that the temporal overlay of neuronal activity and therefore the cooperativity between R2 neurons is the essential factor for generating compound oscillations, which determine the internal sleep drive (**Fig. 6**). Disrupting this cooperativity by expressing Mg^2+^-block-deficient NMDAR interfered with multi-unit synchronization, directly altering the animals’ sleep drive, sleep quality and stimulus-induced wakening. This allows us to establish a causality for network-specific delta-band oscillations in shaping animal behavior (**Fig. 6**).

We here identify slow-wave oscillations as an electrophysiological correlate for sleep regulation and sensory gating in *Drosophila*. Interestingly, oscillatory activity in *Drosophila* R2 neurons is reminiscent of up- and down states occurring at the level of the mammalian thalamus and cortical networks during deep sleep^34^. The R2 network also seems functionally analogous to the thalamus, as network-specific synchronization of slow-wave activity within the thalamus plays a crucial role in maintaining sleep^20-22^ and sensory gating^35^. Comparable to the potential function of network-specific oscillations during ‘local sleep’ in vertebrates, slow-wave oscillations within the flies’ R2 network may well be involved in the homeostatic regulation of synaptic strength^2,7^. We thus suggest that oscillatory network synchronization may represent an evolutionarily selected ‘optimal’ strategy for sleep regulation as well as for the internal representation of sleepiness. Moreover, our mechanistic framework paves the road for identifying evolutionarily conserved fundamental principles that link slow-wave oscillations as electrophysiological hallmarks of sleep to the neuronal processes underlying memory consolidation.

## Materials and Methods

### Experimental preparation, fly lines and drugs

Flies were reared on standard cornmeal food at 25 °C and 60% humidity under a 12 h light/dark regime. We used the driver lines R88F06-GAL4^36^, R76B06-LexA^14^ and R58H05-Gal4^16^, the effector lines UAS-ArcLight^11^, lexAop-CsChrimson, the UAS-RNAi lines^37^ nACHRα1, nACHRα5, nACHRα6, nACHRα7, all obtained from the Bloomington stock center. We also used flies expressing UAS-RNAi against GluCl from the VDRC. UAS-dNR1 wildtype and UAS-dNR1 N631Q (NMDAR^Mg-/-^) were obtained from Dr. Minoru Saitoe^28^ from Tokyo Metropolitan Institute of Medical Science. UAS-dsNR1 and UAS-dsNR2 RNAi lines were obtained from Dr. Chia-Lin Wu from Tsing Hua University.

Whole-brain explant dissections and fly *in vivo* preparation were performed as previously described^11,38^. Low (5 mM) Mg^2+^ external solution consisted of (in mM) 90 NaCl, 3 KCl, 1.5 CaCl_2_, 5 MgCl_2_, 1 NaH_2_PO_4_, 10 glucose, 10 sucrose, 8 trehalose, 5 TES and 26 NaHCO_3_. High (20 mM) Mg^2+^ external solution consisted of (in mM): 70 NaCl, 3 KCl, 1.5 CaCl_2_, 20 MgCl_2_, 1 NaH_2_PO_4_, 10 glucose, 10 sucrose, 8 trehalose, 5 TES and 26 NaHCO_3_. Mg^2+^ concentrations were adjusted accordingly for titration experiments. External solution was adjusted to a pH of 7.4, with an osmolarity of 280 mmol/kg. For focal acetylcholine application, 5-6 consecutive puffs of 1 mM acetylcholine were applied with 1 s breaks consecutively. MK-801 was used at 200 μM (dissolved in 50% EtOH, 2% EtOH in imaging saline). TTX was used at 2 μM (Cayman Chemical) and APV at 100-200 μM (Sigma-Aldrich).

### Voltage imaging and optogenetics

For all experiments 3-10 d old female flies were tested at ZT (Zeitgeber time) 8-16. For optogenetic experiments all-trans retinal (Sigma-Aldrich) was dissolved in 95 % ethanol as a 50 mM stock. 1-3 d female flies were collected after eclosion and transferred to fly food containing 400 μM of the stock solution 2-4 days before imaging.

Imaging was performed on an Olympus BX51WI microscope using a Plan Apochromat 40×, numerical aperture 0.8, water-immersion objective (Olympus, Japan). ArcLight was excited at 470 nm using a Lumencor Spectra X-Light engine LED system. LED power was adjusted for each recording individually to make sure that fluorescent images were not saturated. For optogenetic experiments, CsChrimson was excited at 640 nm. The objective C-mount image was projected onto an Andor iXon-888 camera controlled by Andor Solis software. *Ex vivo* imaging of single R2 neurons and compound activity was mostly performed at a frame rate of 250 Hz. In vivo recordings and recordings in Fig.3 and 4 were performed at frame rates of approximately 70 Hz.

The relative fluorescence change was calculated using the formula: ΔF/F %=[(F_t_-F_b_)/F_b_] * 100. F_t_ is the fluorescence intensity at any given time point t and F_b_ is the baseline fluorescence intensity determined by the applied fitting algorithm at each time point t. In all figures polynomial fitting was applied to compensate for photobleaching. All optical recordings presented here were smoothed using a Savitzky-Golay filter (factor 8-10).

For power spectrum analysis we performed a Fourier-transformation on the unsmoothed relative changes in fluorescence. (The power therefore directly reflects the amplitude of relative changes in fluorescence). The unit is indicated as power = [(ΔF/F)^2^/Hz]*100. For correlation analysis we first generated instantaneous amplitudes ^39^ from the relative changes in fluorescence. Via cross correlation between instantaneous amplitudes of different R2 neurons we obtained a cross correlogramm and main phase lags. The correlation coefficient represents the significance of the main phase lag compared to the phase lag distribution and was calculated using the formula: Coef (X, Y) = Covariance(X,Y)/sqrt(Variance (X)*Variance (Y)), whereas X and Y represent the instantaneous amplitudes of 2 different cells.

### Patch-clamp recordings

*Ex vivo* and *in vivo* whole-cell patch-clamp recordings from R2 neurons were performed as reported elsewhere ^15,40,41^. Identification of R2 neurons was based on GFP expression. External saline was used as described above. Patch pipettes (7-10 MΩ) were filled with internal saline containing (mM): 135 K-aspartate, 10 HEPES, 1 EGTA, 1 KCl, 4 MgATP, 0.5 Na3GTP. Internal solution was adjusted to a pH of 7.2, with an osmolarity of 265 mmol/kg. All signals were digitized at 10 kHz and filtered at 5 kHz. All the analysis was performed with Clampfit 10.7 and MATLAB.

### Hemolymph recordings

Double-barreled ion-sensitive microelectrodes for recording changes in [Mg^2+^]_e_ and [Ca^2+^]_e_ were inserted into the head hemolymph of immobilized tethered flies. The reference barrel was filled with 154 mM NaCl solution, the ion-sensitive barrel with the ionophore cocktail (Magnesium Ionophore II 63083; Calcium Ionophore I 21048; both from Sigma-Aldrich) and 100 mM MgCl_2_ or 100 mM CaCl_2_ respectively. The ion-sensitive microelectrodes were calibrated prior to measurements in the specimen. As magnesium ionophore cocktails also measures Ca^2+^ with the same sensitivity as Mg^2+^, we subtracted the [Ca^2+^]_e_ to the measured [Mg^2+^]_e_.

### Sleep, Sleep Deprivation and Light Stimulus

Flies were isogenized for at least 6 generations. For sleep measurements, single 5- to 6-day-old female flies were loaded into glass tubes (5 mm diameter and 65 mm length) containing 5% sucrose and 2% agar. The activity and sleep of single flies were measured using *Drosophila* Activity Monitors (DAM2) from Trikinetics Inc. (Waltham, MA) at 25 °C under a 12/12 light-dark cycle. Activity counts for each fly were collected every minute for at least 5 days, the data from the first 36 h were excluded due to the entrainment/habitation to new environment. A period of quiescence without activity counts equal to or longer than 5 min was identified as sleep ^5^. Sleep architectures were calculated using Sleep and Circadian Analysis MATLAB Program (SCAMP).

For sleep deprivation, DAM2 monitors were attached to a Vortexer Mounting Plate (Trikinetics) on an Analog Multi-Tube Vortexer controlled by Trikinetics acquisition software. A pulse of mechanical stimulus lasts for 1.2 s and was applied randomly every 20 s. The speed of the vortexer was gradually increased until all flies were sleep deprived during 12 h dark period. For light-induced waking, flies were fully sleep deprived from ZT12 to ZT21, prior to a 10 min light stimulus at ZT22. Duration of the light stimulus was set, so that approximately 50% of control flies were woken. To identify whether a fly was woken or not, we manually examined 3 periods of activity of single flies: 1) 5 min *before* light stimulus, 2) 10 min *during* light stimulus, and 3) 5 min *after* light stimulus. Flies that showed activity *before* the stimulus were excluded. Only flies that were quiescent before the stimulus but showed activity *during* the stimulus were considered awake.

Two-photon images used for schematic overviews were acquired on a Nikon multiphoton microscope at the AMBIO core facility of the Charité.

Statistic tests used and statistic parameters are indicated in the figure legends.

## Supplementary Figures

**Fig. S1:**
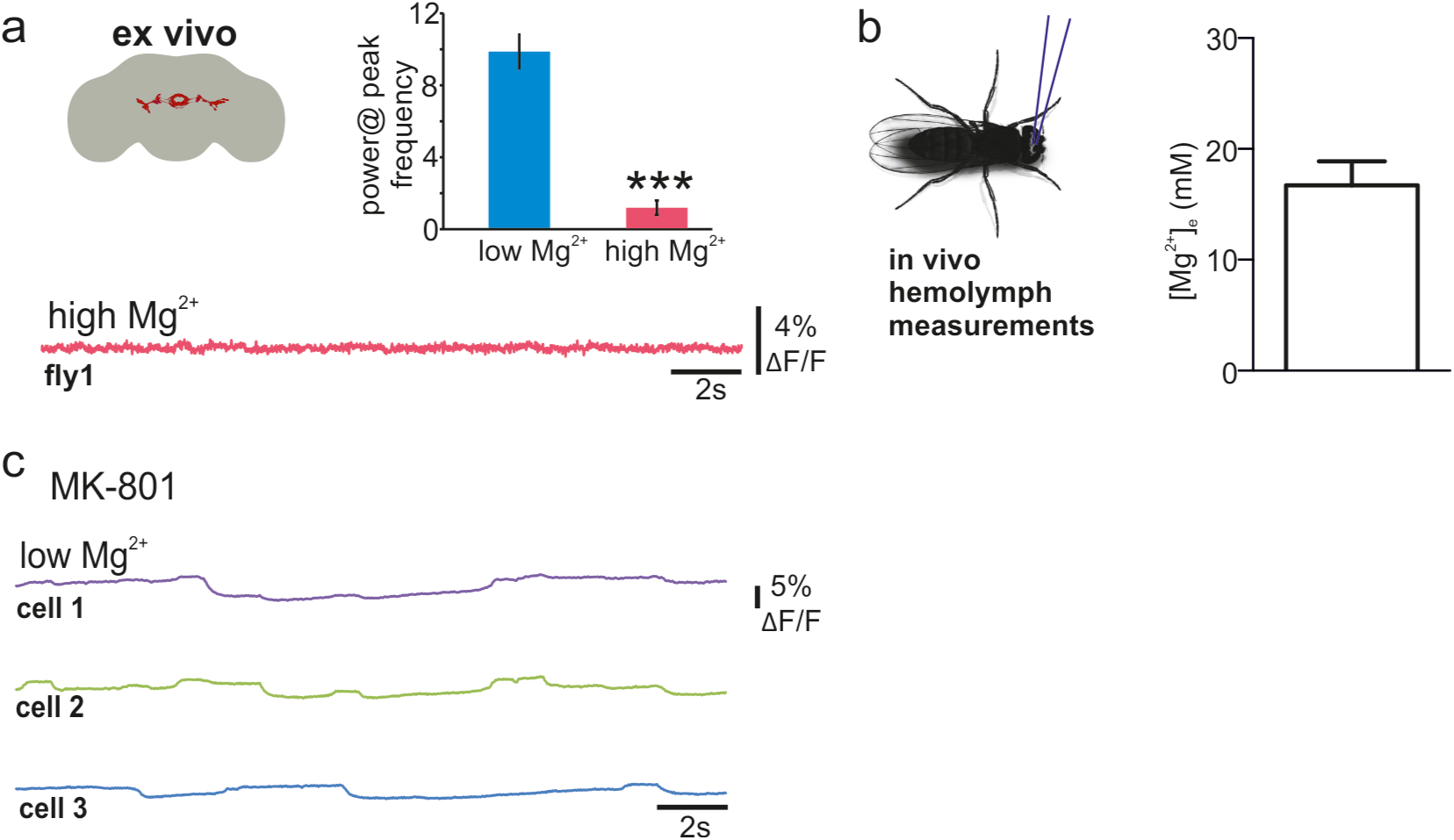
*Ex vivo* compound oscillations and Mg^2+^ concentration in hemolymph. **a,** Compound oscillations are strongly reduced *ex vivo* but reinstated in low external Mg^2+^ (N=7-14 from 7 flies; Mann-Whitney Test, p***<0.001). **b,** Extracellular Mg^2+^ was measured using a Mg^2+^-sensitive electrode inserted into the fly brain. Concentrations measured revealed high Mg^2+^ concentrations (N=19 from 19 flies). **c,** MK-801 abolishes compound oscillations. Bar graphs: mean± sem.

**Fig. S2:**
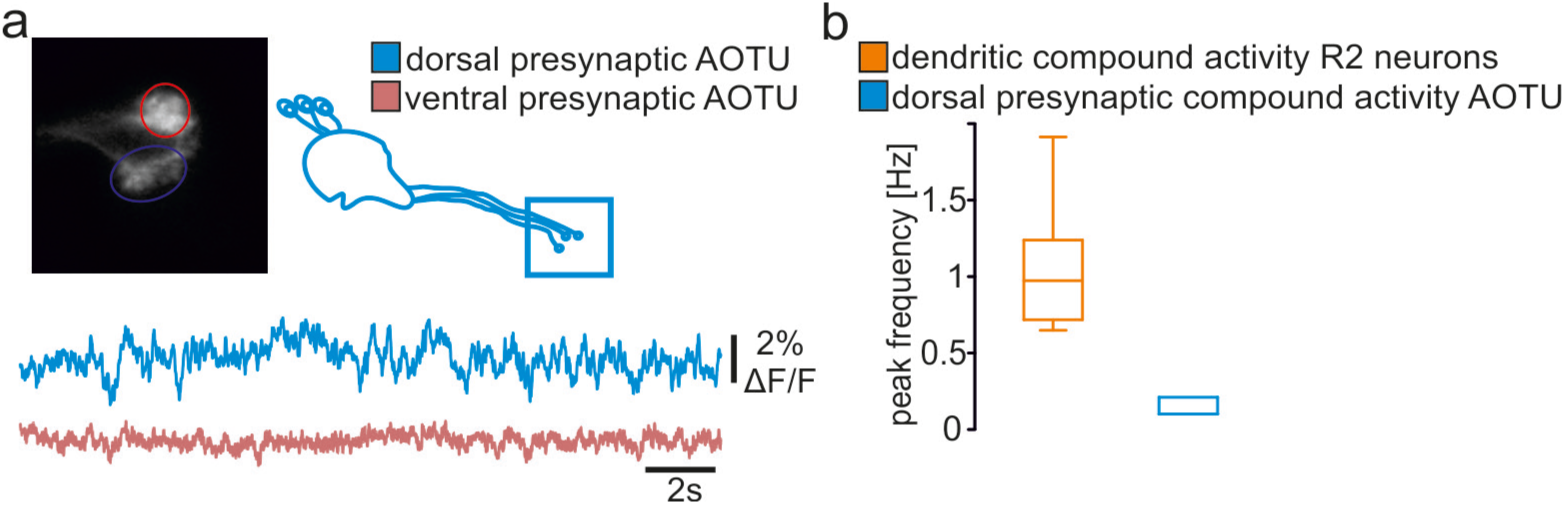
No oscillatory activity in AOTU. **a-b,** No oscillatory activity in dorsal and ventral AOTU *ex vivo* in low (5 mM) Mg^2+^. (N=8-14 from at least 8 flies).

**Fig. S3:**
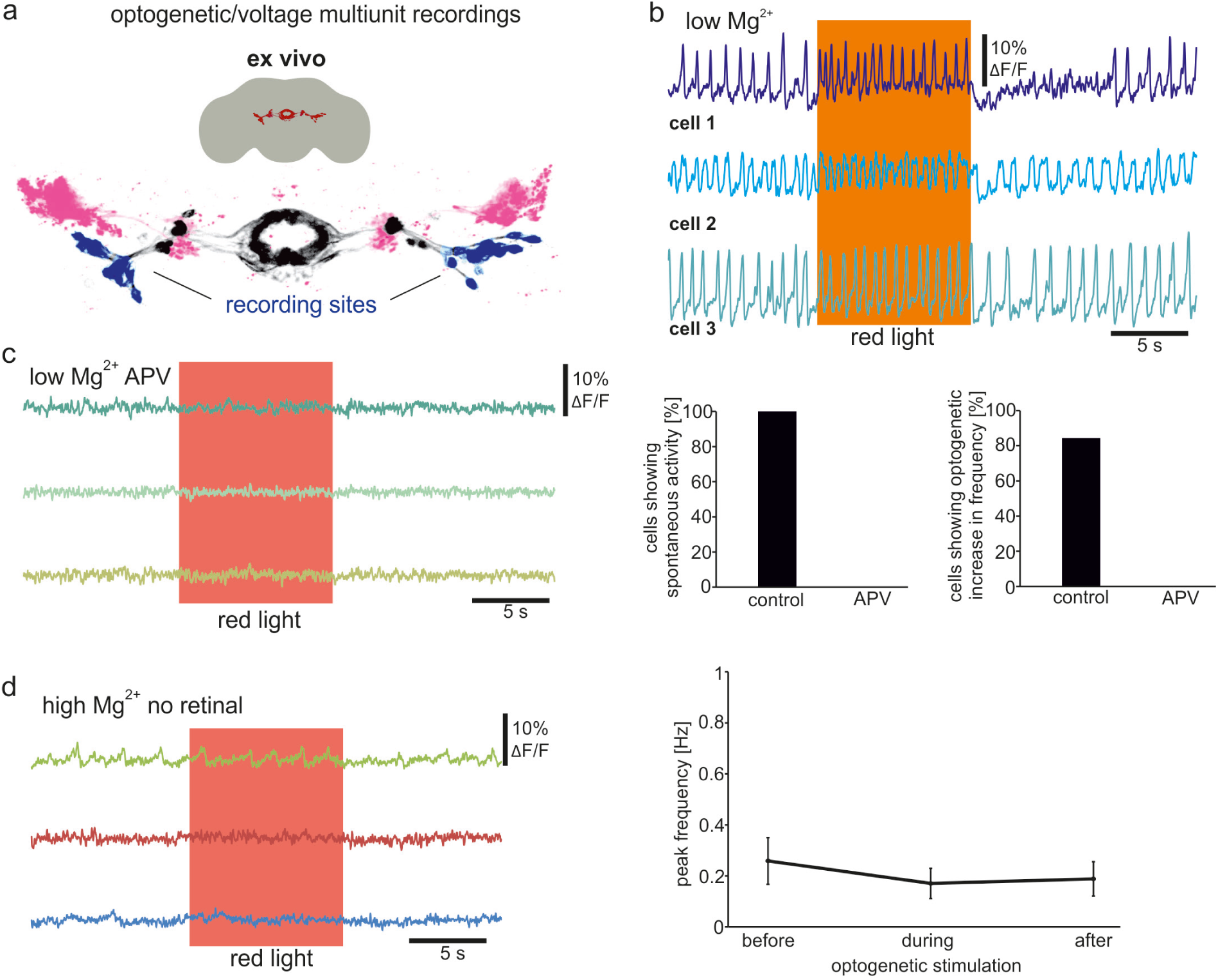
Optogenetically-induced activity in R2 neurons. **a-b,** Optogenetic activation of AOTU induces increased frequency of single-unit activity in low [Mg^2+^]_e_. **c,** In the presence of APV (200 μM), optogenetic stimulation of the AOTU does not induce activity (N=22 from 4 flies). **d,** Peak frequencies in control animals that were not fed with retinal-supplemented food match those in non-optogenetic experiments. No activity change is induced by LED exposure (N=17 from 3 flies).

**Fig. S4:**
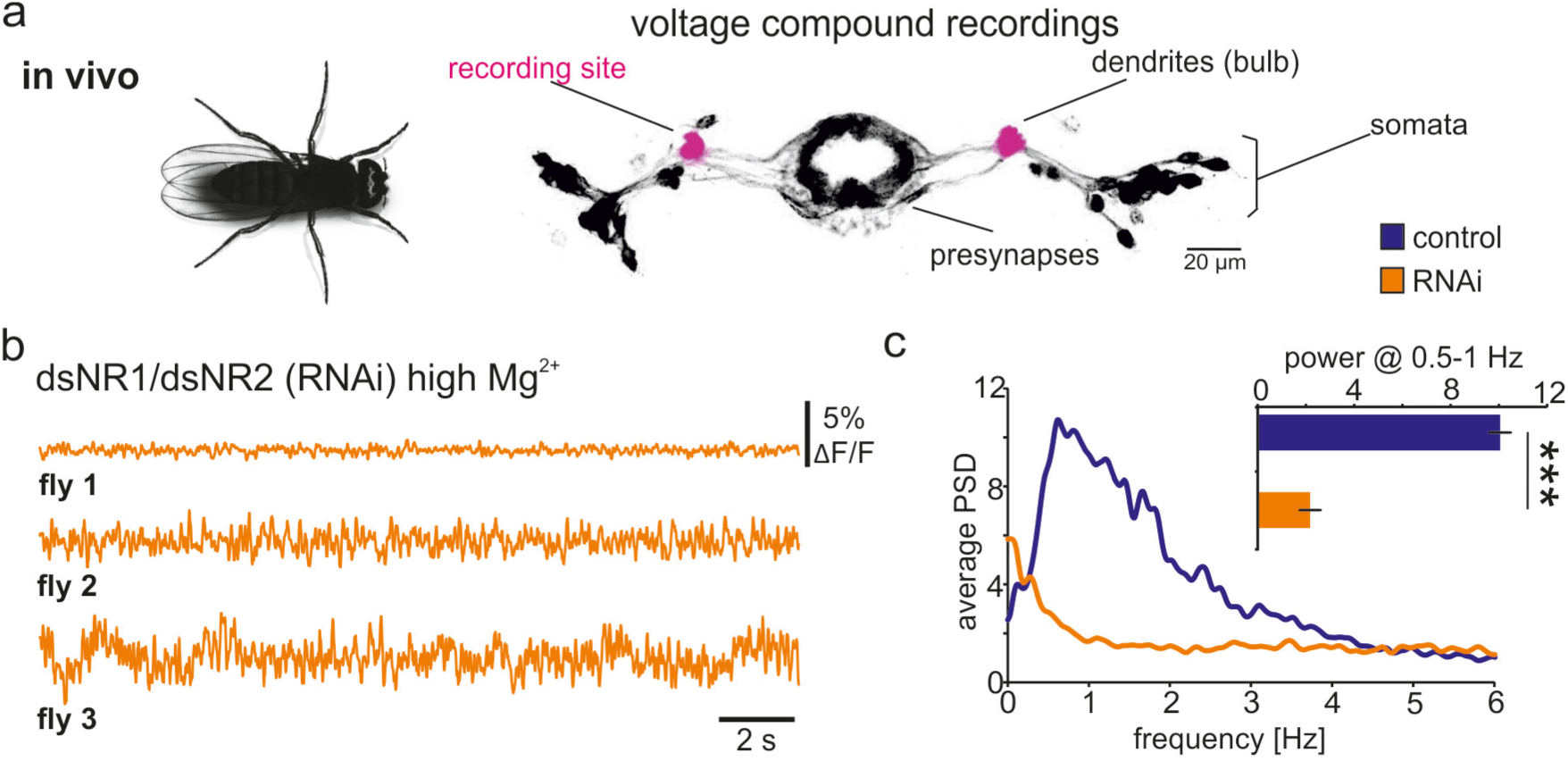
Knock-down of NMDARs *in vivo abolishes compound delta oscillations*. **a-b,** R2 network schematic indicating *in vivo* recording sites. Dendritic compound activity of R2 neurons of flies expressing RNAi against NMDAR S1 and NMDAR S2 (dsNR1/dsNR2) at high [Mg^2+^]_e_. **c,** Power spectrum, NMDAR knockdown abolishes delta oscillations (N=8-10 ; One-way ANOVA followed by, p***<0.001). Bar graphs: mean± sem.

